# The variability of goat microRNA genes is strongly shaped by functional constraints

**DOI:** 10.1101/2025.09.30.679575

**Authors:** Emilio Mármol-Sánchez, María Gracia Luigi-Sierra, Philippe Bardou, Carole Charlier, Licia Colli, Tainã Figueiredo Cardoso, Paola Crepaldi, Arianna Bionda, Marco Milanesi, Barbara Lazzari, VarGoats Consortium, Gwenola Tosser-Klopp, Marcel Amills

**Author notes:** **Corresponding authors:** Emilio Mármol-Sánchez, Marcel Amills.

## Abstract

**Background:** MicroRNAs (miRNAs) are a type of small non-coding RNAs involved in the post-transcriptional repression of target mRNA transcripts, and responsible for the fine-tuning of numerous molecular mechanisms regulating cell metabolism. In goats, multiple miRNAs are involved in coordinating the expression of networks of genes with key roles on the phenotypic variation of milk and meat traits. Although a comprehensive set of goat miRNAs has been annotated, their levels of polymorphism have not been characterized yet. Such information would be relevant in order to explore the effects of miRNA variants on phenotypes of economic interest in goats.

**Results:** By using whole-genome sequencing data from 770 domestic goats with African, Asian, and European origins, we have identified polymorphic sites located within miRNA genes as well as in their flanking regions. In doing so, we have found that miRNA polymorphisms are rare (median alternative allele frequency of 0.46%) and that the distribution of polymorphic sites within and around miRNA loci is uneven. Remarkably, the stem, loop and neighbouring regulatory regions of precursor miRNA hairpins show a significantly higher polymorphism density compared to the miRNA seed, which determines the binding affinity to target mRNAs. Moreover, we have detected a differential segregation of miRNA variants across and within continental regions, with an enriched segregation of putatively high impact polymorphisms, i.e. those located in the seed and other biologically relevant regions of miRNA genes, in isolated goat populations with a low census and elevated content of runs of homozygosity.

**Conclusion:** Goat miRNA genes display low levels of variation particularly in the seed region, likely due to the action of strong purifying selection removing mutations with potential effects on gene regulatory networks linked to miRNA function. Moreover, miRNA polymorphisms tend to be more abundant in goat breeds with high levels of homozygosity, likely because purifying selection is less efficient in populations of limited size. The information provided in the current work could be useful to identify miRNA polymorphisms contributing to phenotypic variation through the disruption of gene regulatory networks in domestic goats, as well as to assess their potential impact on adaptation and fitness.

## Introduction

MicroRNAs (miRNAs) are a type of evolutionarily conserved noncoding regulatory transcripts that fine-tune the expression of hundreds of messenger RNAs (mRNAs) by binding to partially or completely complementary sequences found in their 3’ untranslated regions (UTR) [1], thus hindering their translation in the ribosomes or promoting their degradation via transcriptional slicing and/or poly(A) tail shortening [2]. In this way, miRNAs are essential regulators modulating targeted gene expression in both physiological and pathological conditions [3].

In humans, miRNA variability is strongly constrained by purifying selection, particularly in regions, such as the seed, with critical roles in miRNA function [4]. Indeed, the presence of variant sites within miRNA genes may alter their ability to bind specific sets of mRNAs, or modify their expression, stability or biogenesis, leading to extensive rewiring of regulatory pathways that sometimes result in deleterious consequences for metabolic fitness [5,6]. In goats, several miRNAs involved in the regulation of milk synthesis and growth have been identified. For instance, there is evidence that prolactin and several miRNAs are reciprocally regulated in the mammary gland of ruminants, and different patterns of miRNA expression have been detected in lactating vs. dry goats [7]. Multiple miRNAs have crucial functions related to mammary lipid metabolism, cell differentiation, proliferation and apoptosis [7], or to muscle growth and intramuscular fat deposition [8]. Altogether, these findings suggest that miRNAs might regulate the expression of protein-coding genes with strong effects on milk and meat yield and composition. In domestic ruminants, thousands of miRNAs from bovine, caprine and ovine species were catalogued in the last update of the RumimiR database in 2022 (https://rumimir.sigenae.org; [9]). In their study, Bourdon et al. discussed the convenience of increasing the reach of RumimiR by adding genetic variants of ruminant miRNAs [9], but this has yet to be implemented. Moreover, an in-depth characterization of miRNA variation would be useful to investigate its effects on meat and milk productive traits.

Recent reports have estimated that there are about six hundred domestic goat breeds with a worldwide distribution [10]. The occurrence of founder effects and genetic bottlenecks, combined with the demographic decline of many autochthonous breeds due to their replacement by cosmopolitan breeds, and the progressive abandonment of rural activities have reduced the genetic variation of multiple goat populations [11]. Bertolini et al. [12,13] used an extensive set of goat breeds with SNP chip genotype data (GoatSNP50 BeadChip) to investigate signatures of selection and patterns of runs of homozygosity (ROH) in domestic goats worldwide [14], and identified low levels of genomic diversity in insular Malagasy goats and Northern European breeds [12,15], as well as local adaptations affecting productive traits, coat color and environmental adaptation [13]. In the VarGoats project (https://www.goatgenome.org/vargoats.html), whole-genome sequences from more than 1,000 goats with a worldwide distribution have been generated or retrieved from public databases [16].

In the current study, we have used the VarGoats resource to analyze the diversity of goat miRNA genes in regions subjected to different functional constraints, as well as to characterize the distribution of miRNA variation across various continental populations. In this regard, a low sharing of miRNA variants between geographically distant regions might imply that those variants are evolutionary young. Finally, we wanted to ascertain whether the frequencies of miRNA variants were different in goat populations with high vs. low genomic ROH coverage. Our hypothesis is that miRNA polymorphisms in functional regions might be more abundant in populations with high levels of homozygosity because the removal of deleterious variants by purifying selection is known to be less effective in bottlenecked populations [17].

## Methods

### Annotation of microRNA loci in the goat genome

We first obtained the annotated miRNA sequences available for the goat species from mirBase [18] and miRCarta [19] databases. Subsequently, we aligned both mature miRNAs and pre-miRNA sequences to the ARS1 goat assembly [20](https://www.ensembl.org/Capra_hircus/) by using the bowtie aligner tool [21] without allowing any mismatches and reporting the best possible alignment up to 20 multi-mapped sequences (*bowtie -v 0 -k1 -m 20 --best --strata*). Alternative isoforms of mature miRNAs (isomiRs) were not included in this study. For the chromosome X, we only considered miRNA loci located in the following unplaced scaffolds: NW_017189516.1, NW_017189517.1, NW_017189518.1, NW_017189540.1, NW_017189551.1 and NW_017193093.1. Both NW_017189516.1 and NW_017189517.1 scaffolds have been previously identified as two continuous regions accounting for ∼85.9% of the domestic goat X chromosome [20]. The remaining short scaffolds were selected as belonging to the X chromosome of the ARS1 assembly [20] based on the presence of annotated X-linked miRNAs across mammals. All annotated pre-miRNA (N = 262) and mature miRNA (N = 427) loci in the ARS1 assembly used in this study are available in Additional file 1: **Table S1**.

### Data acquisition, filtering and preprocessing

A dataset of whole-genome sequence data corresponding to domestic and wild goats has been used in this study [16]. Whole-genome sequence data were obtained from the VarGoats Consortium database (https://www.goatgenome.org/vargoats.html). Additional details about data acquisition, variant calling and initial filtering are reported elsewhere [16]. We only considered domestic goats (*C. hircus*) included in the VarGoats dataset. The VCFtools v0.1.16 software [22] was then used to retain single-nucleotide polymorphisms (SNP) within and around (± 1 Kilobases, Kb) miRNA loci by using our custom miRNA gene annotation as reported above in the ARS1 domestic goat assembly [20]. Insertions and deletions were not considered in this study. We then applied an additional quality filter to only allow miRNA SNPs with sequencing depth (DP) ≥ 4 and genotype quality (GQ) ≥10 using the *--minDP 4 and --minGQ 10* parameters from VCFtools v0.1.16 [22]. Individuals and variant sites with missing rates ≥10% were discarded. Only biallelic sites were considered for further analyses. miRNA SNPs located in scaffolds forming part of chromosome X were transformed to haploid genotypes in male goats using the fixploidy plugin from bcftools v1.21 software [23]. This resulted in a total of 206 miRNA SNPs segregating across 770 domestic goats from African (N = 373), Asian (N = 136), and European (N = 261) origin. These populations were further subdivided according to their geographic intracontinental distribution into Northern African (N = 165), Western African (N = 36), Eastern African (N = 172), Eastern Asian (N = 72), Southern Asian (N = 64), Northern European (N = 26), Western European (N = 211), and Southern European (N = 24) domestic goats. Minimum allele frequencies were calculated with the *--freq* option of VCFtools v0.1.16 [22]. Finally, we created independent genotype files for domestic goats from African, Asian and European origin. Summarized metadata from all the domestic goats (N = 770) considered in this study is available in Additional file 2: **Table S2.**

### Distribution of polymorphisms along microRNA loci in the goat genome

We aimed to investigate the patterns of SNP distribution specifically within miRNA genes and their flanking regions. In this way, we sought to determine whether the amount of diversity differs across miRNA regions subjected to distinct functional constraints due to their biological relevance.

MicroRNAs are first transcribed as a long primary miRNA transcript (pri-miRNA) encompassing the miRNA locus plus additional flanking nucleotides (nt) where processing motifs are typically located [1]. Shortly after being transcribed, the pri-miRNA transcript is processed by the DROSHA-DGCR8 complex in the nucleus to generate a precursor hairpin (pre-miRNA) encompassing two paired stems by imperfect complementarity and an apical loop connecting them. Once processed, the pre-miRNA is then translocated to the cytoplasm where the Dicer protein cleaves the apical loop to produce a ∼22-base pair (bp) miRNA duplex formed by the two initial hairpin stems. This duplex includes both the 5’ prime (5p) and 3’ prime (3p) mature miRNA strands within, each of which of ∼22 nt long, although only one of them is typically functional and loaded to the guide-strand channel of Argonaute (AGO) protein to form the miRISC complex for active mRNA targeting, while the other strand is degraded [1]. The 1^st^ nt from the 5’ sequence end of the guide mature miRNA is not involved the miRNA binding specificity but instead it functions as an anchor to the MID domain of AGO to stabilize the miRISC complex. The 2^nd^ to 8^th^ nt of each mature miRNA are the main binding determinants to target mRNAs, known as the miRNA seed. The following 9^th^ to 12^th^ nt are not involved in any functional activity among metazoan miRNAs, and merely serve as a bridge towards the 13^th^ to 18^th^ nt of the mature miRNA, which may provide additional pairing specificity to the seed, thus reinforcing the binding to target mRNAs [1]. Similar to nucleotides 9^th^ to 12^th^, the last 19^th^ to ∼22^nd^ nt are not involved in any biological function. A schematic representation of a metazoan miRNA transcript including all the aforementioned regions is depicted in **Fig. 1**.

**Fig. 1:**
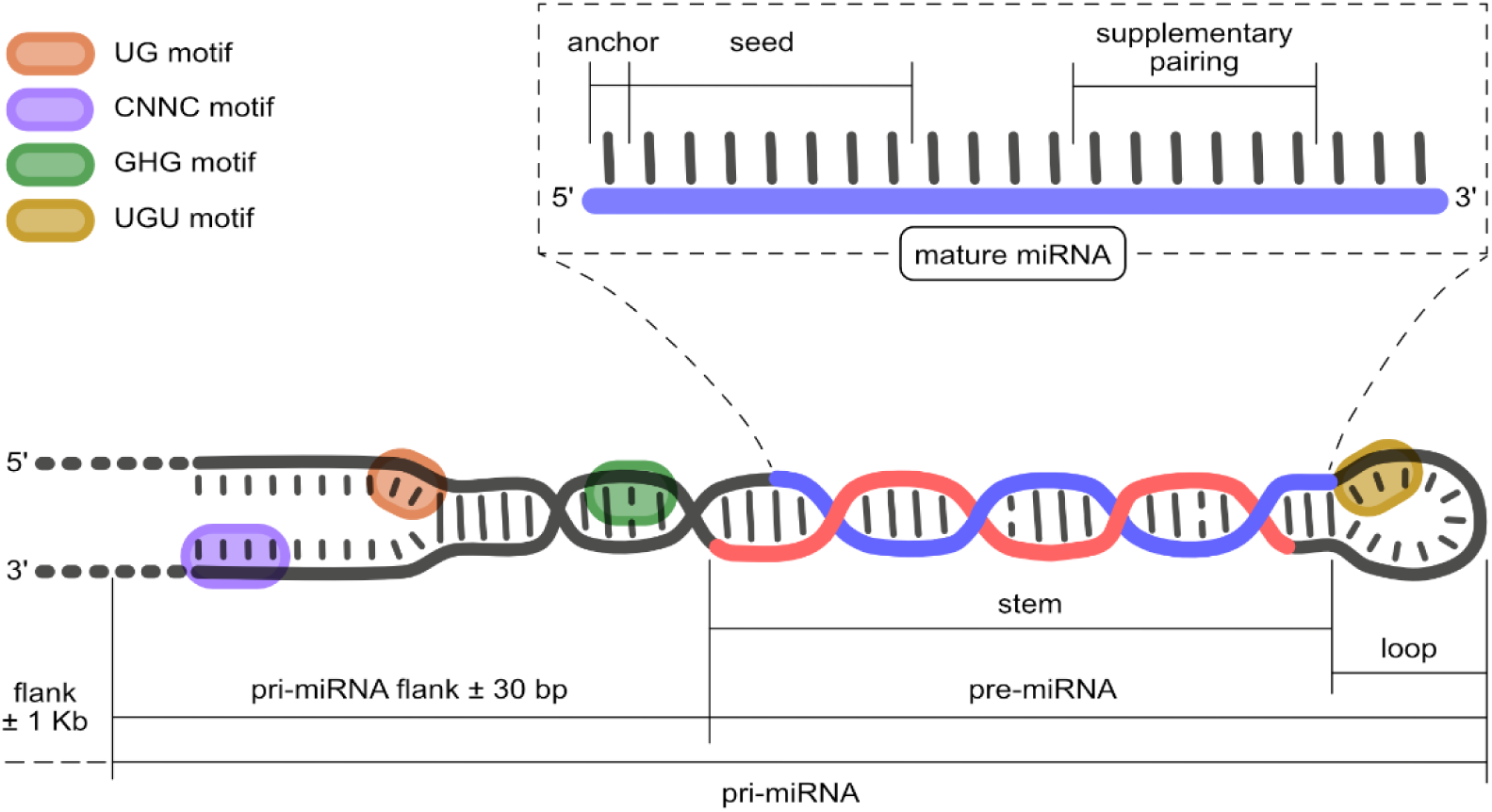
Schematic representation of the microRNA gene regions analyzed in this study. The 5’ (5p) mature miRNA is depicted in blue, while the 3’ (3p) mature miRNA is depicted in red.

We thus established the following regions within and around each miRNA gene: (i) pri-miRNA flank ± 30 bp, defined by the 30 nt immediately surrounding the pre-miRNA locus; (ii) pre-miRNA stem/loop, defined by the pre-miRNA hairpin stem and the apical loop, excluding either the 5p or 3p mature miRNA strands, if annotated. For miRNA genes where only one mature miRNA strand was annotated (either the 5p or 3p), the complementary strand was included as part of the stem/loop region; (iii) mature miRNA, defined by the 427 mature miRNAs annotated in the goat genome considered in this study, and excluding the seed region; (iv) miRNA seed, defined by the 2^nd^ to 8^th^ nt from each annotated mature miRNA; and (v) flanking ± 1 Kb, defined by genomic regions located 1 Kb upstream and downstream of each pri-miRNA locus identified by us (N = 262). Moreover, we defined two additional subregions within each mature miRNA: anchor (1^st^ nt from the 5’ sequence end) and supplementary pairing (13^th^ to 18^th^ nt).

To determine the SNP density within each of the regions defined above, we calculated their corresponding additive length per chromosome, excluding autosomes 6, 9, and 17, where no miRNAs were annotated. Then, the miRNA SNP density (*D*) of each one of these regions was calculated as follows:

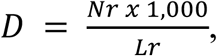

where *Nr* is the number of detected SNPs per region and chromosome, and *Lr* is the nucleotide length of each defined region per chromosome. The resulting density metric was then transformed to the number of SNPs per Kb by multiplying *Nr* values per 1,000 in the aforementioned formula. To assess the existence of statistically significant differences among *D* values across miRNA regions, we used a pairwise comparison across miRNA regions using the Mann-Whitney U non-parametric test [24] with the *pairwise.wilcox.test* function in R v4.4.2 [25]. Density estimates across different chromosomes for a given miRNA region were treated as independent replicates. The resulting *P*-value estimates were adjusted for multiple testing with the Bonferroni method [26].

### Principal Component Analysis

Population structure based on miRNA SNPs was determined by using PLINK v1.9 software [27], and including variants within and around ± 30 bp miRNA loci as described above. Subsequently, the *--pca* option was used to carry out Principal Component Analysis (PCA) obtaining the first 20 principal eigenvectors. Additional chromosomes present in the goat assembly were taken into account with the *--chr-set 37 --allow-extra-chr* options. After PCA, the resulting eigenvalue vectors were used to estimate the percentage of variance (V) explained by each of the first 20 principal components using the following formula:

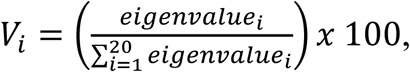

where the variance *V* explained by each *i* top 20 principal components is expressed as the ratio between each *i eigenvalue* and the cumulative sum of the eigenvalues of the 20 eigenvectors, then multiplied by 100 to express a percentage value. PCA was computed including all 770 domestic goats analyzed in this study, as well as independently for each of the specific subsets of goats from Africa (N = 373), Asia (N = 136) and Europe (N = 261).

### Distribution of miRNA variation in goat breeds with high and low ROH coverage

Additionally, we aimed to determine whether domestic goat breeds with high or low levels of genomic homozygosity showed differential segregation patterns of miRNA SNPs potentially affecting miRNA expression, processing and maturation, or target binding specificity. In this way, we used estimates of the average fraction of the goat genomes containing ROHs for multiple domestic goat breeds reported by Bertolini et al. [12]. We then established a maximum threshold of 5% for goat breeds classified as having low genomic ROH content, and a minimum threshold of 10% for goat breeds with high genomic ROH content.

A total of 74 goats with high ROH content (14–35%) were present in our dataset and they belonged to the following breeds or regional subdivisions: Bari (N = 4), Dedza (N = 3), Diana (N = 7), Girgentana (N = 4), Kachan (N = 4), Kamori (N = 5), Landin (N = 6), Landrace (N = 15), Maltese (N = 2), Menabe (N = 8), Old Irish (N = 4), Palmera (N = 2), Pateri (N = 2), and Sofia (N = 8). On the other hand, a total of 77 goats with low ROH content (2–4%) belonged to the following breeds: Abergelle (N = 6), Bermeya (N = 5), Galla (N = 7), Gogo (N = 7), Gumez (N = 4), Maasai (N = 8), Maure (N = 4), Peulh (N = 4), Rossa Mediterranea (N = 3), Sonjo (N = 7), Soudanaise (N = 8), Targui (N = 3), Tunisian (N = 3), and Woyito Guji (N = 8). Thus, a total of 151 goat samples were considered in these analyses. To strengthen allele frequency estimates, we grouped the breeds onto geographic subregions to ensure that no group had less than 5 goat samples. More specifically, we defined the following geographic groups for the high genomic ROH content: Malagasy (N = 23, including Diana, Menabe, and Sofia goats), Mediterranean (N = 6, including Girgentana, and Maltese goats), African (N = 11, including Palmera, Dedza, and Landing goats), North European (N = 19, including Landrace and Old Irish goats), and Pakistani (N = 15, including Bari, Kachan, Kamori and Pateri goats). For goat breeds with low genomic ROH content we defined the following geographic groups: Mediterranean (N = 19, including Bermeya, Rossa Mediterranea, Soudanaise, and Tunisian goats), Ethiopian (N = 18, including Abergelle, Gumez and Woyito Guji goats), East African (N = 29, including Galla, Gogo, Maasai and Sonjo goats), and West African (N = 11, including Maure, Peulh and Targui goats). A summary of the aforementioned breeds, their estimated ROH values, and assigned geographic groups is available in Additional file 3: **Table S3**.

### Weighted allele frequencies

To ensure an unbiased comparison between alternative allele frequencies across groups of goat breeds differing in sample size, we incorporated a weighting factor *w* to account for differences in population size for each group defined above. In this way, a within-group weighted alternative allele frequency (*wFrq*) was computed for each *j* miRNA polymorphic site expressed as:

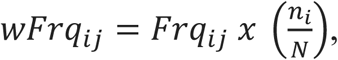

where *Frq_ij_* is the alternative allele frequency for each *j* miRNA polymorphism calculated independently for each *i* group, *N* is the population size considered for each goat breed with high or low genomic ROH content, and *n* is the number of goat individuals for each *i* breed. Subsequently, to improve the visualization of the data, weighted allele frequencies for each miRNA SNP were standardized (z-score transformation) and grouped based on hierarchical clustering over Euclidean distances calculated with the *dist* and *hclust* functions in R v4.4.2. [25].

### Allele frequency change

When comparing the alternative allele frequencies among goats from breeds with high genomic ROH content vs. breeds with low genomic ROH content, we calculated the corresponding frequency change as the differential between allele frequencies from the high and low groups, respectively, expressed as:

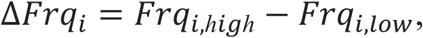

where *Frq* is the corresponding alternative allele frequency for each *i* miRNA SNP among breeds with *high* and *low* ROH content according to Bertolini et al. [12].

### Polymorphism prioritization

To investigate the segregation patterns of miRNA SNPs contributing the most to differential clustering patterns after PCA for goats with high and low ROH content as described above, we obtained genotype information for each individual and each selected SNP within or nearby miRNA loci (up to ± 30 bp upstream and downstream). Homozygous individuals for the reference alleles were assigned a genotype value of zero, while individuals with heterozygous or homozygous genotypes for the alternative alleles were assigned genotype values of one and two, respectively. For SNPs located in scaffolds assigned to the X chromosome in male goats, homozygous alleles for the reference allele were assigned a value of zero, while homozygous alleles for the alternative allele were assigned a value of one. Prior to standardization of genotype values (z-score transformation), we imputed missing genotypes for each polymorphic miRNA SNP by assigning to them the mean genotype value across all individual goats considered in these analyses. We then calculated the contribution or loading of each polymorphic site to the PCA linear embedding by multiplying the standardized imputed genotype matrix by the top 20 principal component eigenvectors for each sample. Varimax rotation was then applied to loading values in order to maximize the variance explained for each SNP within each considered component. The varimax function within R v4.4.2 [25] was used for such calculation. The contribution of each polymorphism to clustering linear patterns observed after PCA was then estimated by computing a weighted score (W) defined as:

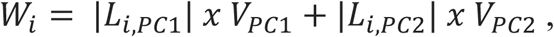

where *L* denotes the absolute values of varimax-rotated loadings for each *i* miRNA SNP over the first (PC1) and second (PC2) principal components, and *V* is the percentage of variance explained by each PC1 and PC2 components, respectively.

Finally, the miRNA SNPs that contributed the most to PCA discrimination between goats with high and low ROH content were identified using a two-component Gaussian mixture model with variable variance across components [28]. Variant sites with weighted scores (W) assigned to the component with the higher mean value and with a posterior probability >95% of belonging to the assigned component were selected as the top discriminating miRNA SNPs.

### Functional enrichment

To evaluate whether a specific miRNA functional region was significantly enriched among the full set of miRNA SNPs, and whether variants with higher alternative allele frequencies in goat breeds with high ROH content compared to those with low ROH content [12] were also enriched among the top most discriminating miRNA SNPs, we performed a one-tailed Fisher’s exact test using the fisher.test function in R v4.4.2 [25]. Statistical significance was defined at a nominal *P*-value < 0.05.

## Results

### Single nucleotide polymorphism density within miRNA loci is strongly biased across functional regions

After mapping goat miRNAs to the ARS1 genome assembly, a total of 427 mature miRNAs were successfully annotated within 262 different pre-miRNA loci. The detection of variant sites located across each miRNA gene, as well as in their flanking regions (± 30 bp) was carried out by using a collection of whole-genome sequences from 770 domestic goats (*C. hircus*) from African, Asian and European origin belonging to the VarGoats dataset after quality filtering. A total of 206 segregating variants, located within 123 distinct miRNA loci, were identified. From these, 74 out of 123 miRNAs (60.16%) harbored just one variant site, 27 miRNAs (21.95%) showed two, 14 miRNAs (11.38%) showed three, and eight miRNAs (6.50%) harbored up to six variants (Additional file 4: **Table S4**). Conversely, out of the 262 annotated miRNA loci, 139 (53.05%) lacked any variant within their precursor (pre-miRNA) or the ± 30 bp flanking regions, so they were classified as monomorphic. A total of 23 out of the 206 SNPs (11.16%) were only found in either homozygous or heterozygous state in one single domestic goat individual, and thus were classified as singletons (Additional file 4: **Table S4**). Interestingly, seven of these SNPs were predicted to modify miRNA processing motifs (see **Fig. 1** for a schematic representation of their location within the miRNA hairpin and their flanking regions). This finding was deemed as a significant enrichment (*P*-value = 8.4E-03) for extremely rare miRNA SNPs (i.e. singletons) putatively altering miRNA processing motifs compared to other variant sites found in more than one goat individual.

Among the 206 miRNA SNPs detected, only three (1.45%) were located in the seed region (2^nd^ to 8^th^ nt) of chi-miR-1197, chi-miR-425-3p and chi-miR-1468-3p mature miRNA sequences, respectively (Additional file 4: **Table S4**). Moreover, eighteen variants (8.74%) were also located within mature miRNA regions, but outside the seed. From these, one variant was located in the anchor region (1^st^ nt of the mature miRNA), four were located in the supplementary pairing region (13^th^ to 18^th^ nt), and the remaining thirteen were located in regions without any known functional role (i.e. 9^th^ to 12^th^, and 19^th^ to the 3’ end of the mature miRNA sequence; Additional file 4: **Table S4**). Besides, 51 variants (24.76%) mapped to the pre-miRNA precursor region, but outside the annotated mature and seed miRNA regions. From these, 24 were found in the hairpin stem and 27 were located in the apical loop. The majority of miRNA SNPs (N = 134, 65.05%) were found in the 30 bp immediately upstream or downstream to the pre-miRNA hairpin boundaries, also known as the pri-miRNA flanking region, where miRNA processing motifs are typically located (see **Fig. 1**, and Additional file 4: **Table S4**). When considering miRNA SNPs beyond the pri-miRNA flanks, up to 1 Kb upstream and downstream, a total of 7,372 variants were found.

Overall, the median value of the alternative allele frequencies observed for these 206 miRNA SNPs across all analyzed domestic goats (N = 770) was 0.46% (**Fig. 2a**), meaning that the vast majority of these SNPs were found only in a very small number of goats. However, certain miRNA SNPs showed higher values, with 61 out of the 206 (23.5%) having alternative allele frequencies above 2% (mean alternative allele frequency = 4.22%, see Additional file 4: **Table S4**). None of these variant sites with relatively high frequencies were located in the miRNA seed, but instead they were predominantly found in pri-miRNA flanking regions or in the pre-miRNA stems and apical loops. Only one miRNA SNP with an alternative allele frequency >2% was found in the mature miRNA region, but it resided outside the seed (21:65224784, rs666792599, located at chi-miR-377-3p, alternative allele frequency = 2.55%; see Additional file 4: **Table S4**).

**Fig. 2:**
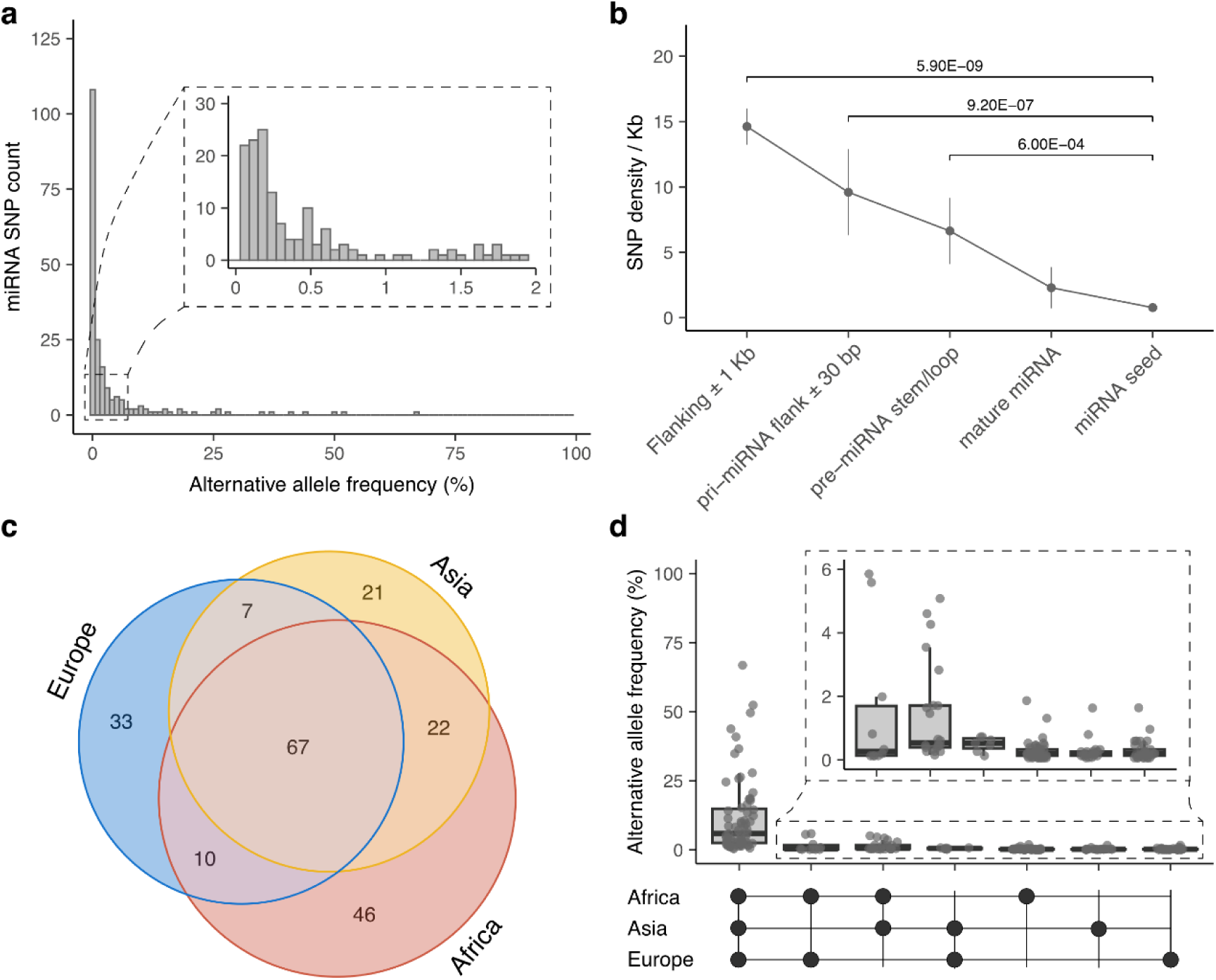
Distribution of miRNA polymorphisms in domestic goat populations. (**a**) Alternative allele frequency (%) distribution of miRNA SNPs (N = 206) detected in the population of 770 domestic goats from the VarGoats project analyzed in this study. (**b**) miRNA SNP density (measured as the number of polymorphic sites per Kilo base-pair, Kb) across miRNA regions defined as: i) Flanking regions at ± 1 Kb upstream and downstream the pri-miRNA hairpin boundaries; ii) pri-miRNA flank ± 30 bp, defined by the 30 nucleotides immediately surrounding the pre-miRNA upstream and downstream boundaries; (iii) pre-miRNA stem/loop, defined by regions within the miRNA gene encompassing the hairpin stem and the apical loop without overlapping annotated mature miRNAs; (iv) mature miRNA, defined by the annotated mature miRNA loci considered in this study (N = 427); and (v) miRNA seed, defined by the 2^nd^ to 8^th^ nucleotide from each annotated mature miRNAs. Error bars for each miRNA region represent the standard error considering all SNP density measured across all annotated miRNA loci wherever a SNP site was detected. (**c**) Venn diagram depicting the sharing of miRNA SNPs (N = 206) by African, Asian, or European domestic goats. (**d**) Boxplots depicting the alternative allele frequencies (%) of miRNA SNPs unique to or shared among the continental goat populations under study, with intersections represented by an UpSet matrix (bottom).

When we compared the estimated SNP density for each of the defined miRNA regions, the scarcity of SNPs in miRNA seeds or mature regions contrasted with the higher SNP density observed for pre-miRNA stems/loops and flanking regions (**Fig. 2b**). Indeed, the SNP density in miRNA seeds (0.8 SNPs/Kb) was ∼3-fold lower than that of mature miRNA regions (2.3 SNPs/Kb) and significantly lower (∼8.5-fold, *P*-value = 6.00E-04) than that of the pre-miRNA stem/loop (6.6 SNPs/Kb, **Fig. 2b**). Similarly, the SNP density obtained for mature miRNA regions was ∼3-fold lower compared to that of pre-miRNA stem/loop though not at significant levels. The SNP density of pri-miRNA flanking regions (± 30 bp, with 9.6 SNPs/Kb) was significantly higher compared to the seed region (*P*-value = 9.20E-07, **Fig. 2b**). Moreover, the flanking regions defined by 1 Kb upstream and downstream miRNA loci showed a mean SNP density of 14.6 SNPs/Kb, which was ∼1.5-fold and ∼2.2-fold significantly higher than the observed density for pre-miRNA and pri-miRNA regions, respectively, as well as ∼6.3-fold and ∼19-fold significantly higher compared to mature miRNA regions and to the miRNA seeds (**Fig. 2b**). Statistical significances derived from all pairwise comparisons of the SNP densities among the miRNA regions analyzed are shown in **Table 1**.

**Table 1.** Pairwise comparison of the SNP densities among all miRNA regions analyzed^a^.

| | Flanking<br>± 1Kb | pri-miRNA flank<br>± 30 bp | pre-miRNA<br>stem/loop | mature<br>miRNA |
| --- | --- | --- | --- | --- |
| pri-miRNA flank ± 30 bp | 1.26E-03 | - | - | - |
| pre-miRNA stem/loop | 7.90E-05 | 1.00E+00 | - | - |
| mature miRNA | 2.60E-08 | 3.40E-04 | 8.76E-02 | - |
| miRNA seed | 5.90E-09 | 9.20E-07 | 6.00E-04 | 4.73E-01 |

<sup>a</sup> $P$ -values are estimated using a Wilcoxon rank-sum test (Mann-Whitney U non-parametric test [24]) and corrected for multiple testing with the Bonferroni method.

### Limited sharing of miRNA polymorphisms across different goat continental populations

The segregation patterns of miRNA SNPs located up to ± 30 bp flanking the pre-miRNA boundaries showed that 67 out of the 206 SNPs (32.52%) were shared among African, Asian and European goats (Additional file 5: **Table S5**). miRNA SNPs specifically shared between two of the three continental populations were fewer, with almost half of the identified miRNA SNPs (N = 100) exclusively segregating in either African, Asian or European goats (**Fig. 2c**). African goats showed the highest number of population-specific miRNA SNPs (N = 46), followed by European (N = 33), and Asian goats (N = 21), see Additional file 5: **Table S5**. Interestingly, miRNA SNPs shared among African, Asian and European goat breeds were the ones predominantly showing increased alternative allele frequencies (Additional file 5: **Table S5**, **Fig. 2d**), while population-specific SNPs were overall found in a reduced number of goat individuals. Moreover, two out of the three miRNA SNPs located within the seed region were found in goats with European origin, while the other one was detected in Southern Asian goat populations from Iran and Pakistan, being all of them in a heterozygous state (Additional file 6: **Table S6**).

PCA visualization including all 770 domestic goats analyzed in this study revealed a clearly divergent clustering among African, Asian and European populations (**Fig. 3a**, Additional file 7: **Table S7**). Besides, the analysis of PCA clustering patterns within each of the three continental regions analyzed revealed that goats from Madagascar (including most samples from Androy, Diana, Menabe, Sofia and Sud Ouest regions) showed patterns of miRNA SNP segregation that diverged from those of West and North African goat breeds (**Fig. 3b**). Likewise, goats from Norwegian and Landrace breeds clustered far apart from the main group of European breeds (**Fig. 3c**). With regard to Landrace goats, only those from Finish and Netherlands, but not all of them, clustered apart from the remaining goat individuals (**Fig. 3c**, Additional file 8: **Table S8**). Conversely, Asian breeds showed a more compact clustering, although two goats from Iran were outliers (**Fig. 3d**, Additional file 8: **Table S8**). The corresponding genotype of each miRNA SNP (N = 206) from all domestic goats considered in this study (N = 770) is available in Additional file 9: **Table S9**.

**Fig. 3:**
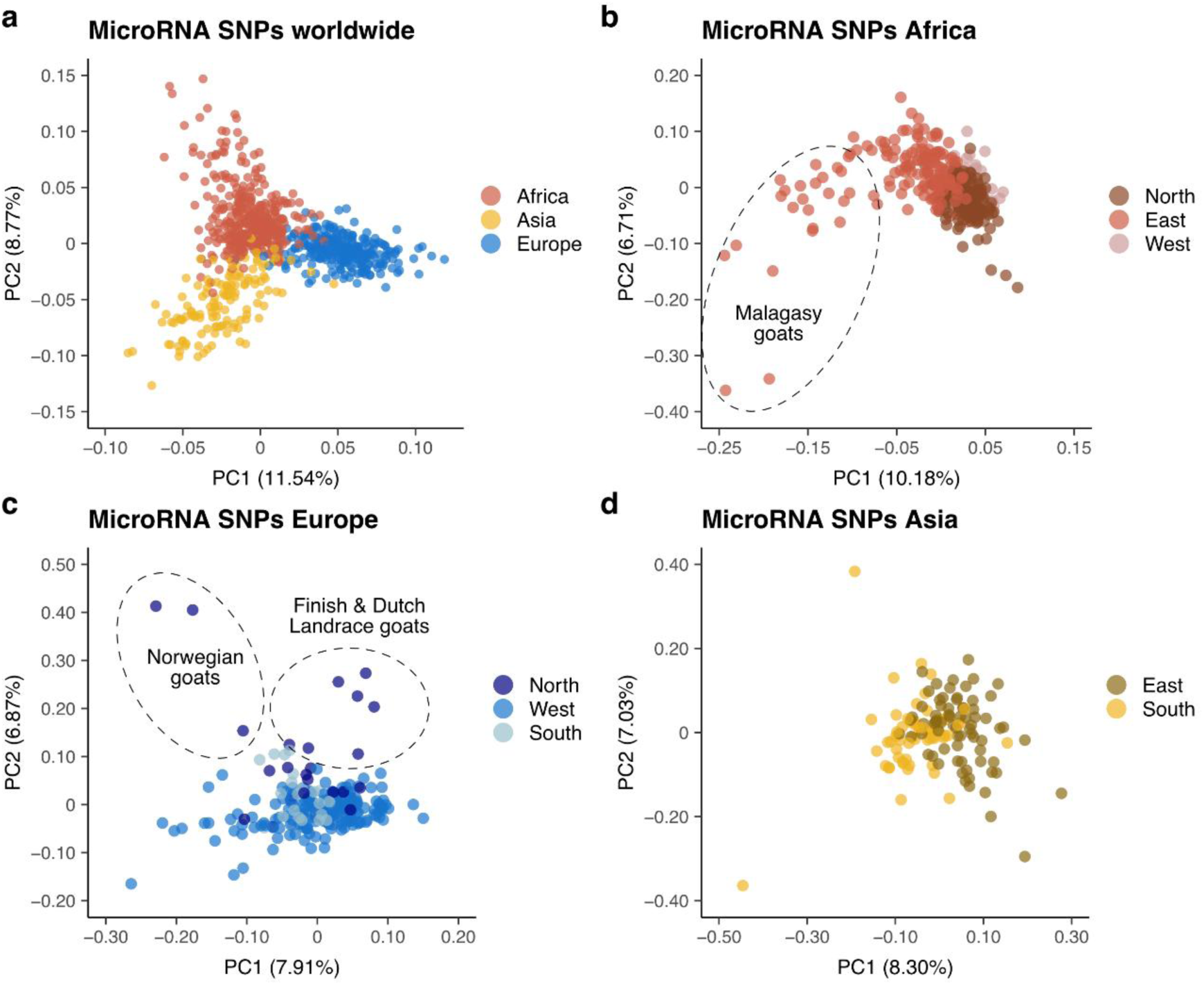
Principal Component Analysis (PCA) of miRNA polymorphisms segregating in domestic goat populations. Each dot in the PCA represents an individual goat sample. (**a**) PCA considering all domestic goats with African, Asian, and European origin from the VarGoats project analyzed in this study (N = 770). (**b**) PCA considering domestic goats from Africa (N = 373). Goats are divided into Northern, Western or Eastern Africa according to their regional geographic origin. (**c**) PCA considering domestic goats from Europe (N = 261). Goats are divided into Northern, Western or Southern Europe according to their regional geographic origin. (**d**) PCA considering domestic goats from Asia (N = 136). Goats are divided into Eastern or Southern Asia according to their regional geographic origin. Principal component eigenvectors for each African, European, and Asian goat population analyzed can be found in Additional files 7 and 8: **Tables S7**-**8**.

### miRNA polymorphisms with putative deleterious effects are more abundant in highly homozygous goat breeds

We aimed to investigate whether goat populations with a high ROH content would have accumulated an elevated burden of miRNA SNPs in functional regions critical for miRNA regulatory function. In order to test this hypothesis, we used a list of goat breeds and their ROH content estimates (expressed as a percentage of their genome covered by ROHs) reported by Bertolini et al. [12]. This included a total of 151 domestic goats that were divided into breeds with high (N = 74 goats, belonging to 14 different breeds, and grouped in 5 geographic regions) and low (N = 77 goats, also belonging to 14 different breeds, but grouped in 4 different geographic regions) ROH content. Goats within the “*high*” group in our dataset had an average ROH content of 21.07% (± 6.79%), while goats in the “*low*” group had an average ROH content of 3.14% (± 0.66%) according to data reported by Bertolini et al. [12] (Additional file 3: **Table S3**). A total of 126 miRNA SNPs were segregating among breeds with high or low ROH contents (Additional file 10: **Table S10**). Sample clustering based on miRNA SNP variation within breeds with high and low ROH contents revealed three groups of goats with high ROH content that showed clearly differentiated segregation patterns when compared to goats with low ROHs (**Fig. 4a**, Additional file 11: **Table S11**). These three groups included Girgentana, Landrace and Old Irish goats with a European descent, African Malagasy goats sampled in Menabe, Sofia and Diana regions, as well as Pakistani goats belonging to the Bari, Pateri, Kamori and Kachan breeds (**Fig. 4a**, Additional file 10: **Table S10**). Among the 126 segregating miRNA SNPs, 83 (65.87%) were located within the pri-miRNA ± 30 bp flanking regions, and 13 of them were predicted to potentially alter miRNA processing motifs (Additional file 12: **Table S12**). Besides, 33 SNPs (26.19%) were located in either the pre-miRNA stem or the hairpin loop, eight (6.35%) were located within mature miRNA sequences, but outside any functionally active region, and two (1.59%) were located in the miRNA seeds of chi-mir-1197-3p (21:65190342) and chi-miR-425-3p (22:51044384, rs666718867), respectively (see Additional file 12: **Table S12**). In accordance with our expectation, 81 out of the 126 SNPs (64.29%) showed increased alternative allele frequencies in breeds with a high ROH content (Additional file 12: **Table S12**).

**Fig. 4:**
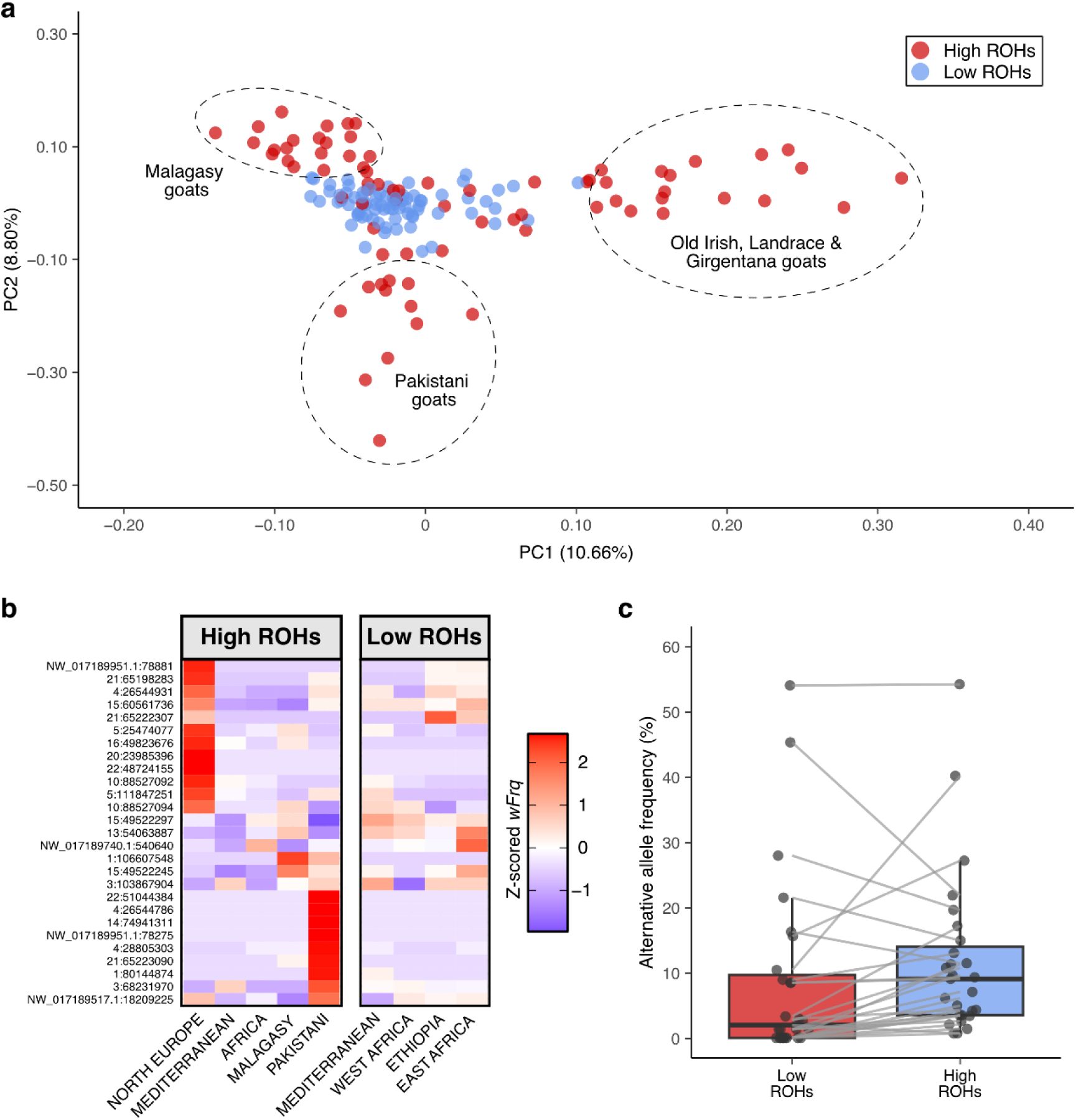
Divergent clustering and allele frequencies in domestic goats with high or low genomic ROH content. (**a**) PCA considering domestic goats from breeds with high or low genomic ROH content according to Bertolini et al. [12] (N = 151). Each dot in the PCA visualization represents an individual goat sample. (**b**) Heatmap representing standardized weighted alternative allele frequencies (*wFrq*) of miRNA SNPs with the top most contribution to the linear clustering detected after PCA (N = 27). Polymorphic sites with higher standardized wFrq values (in red) are predominantly segregating in the corresponding goat group. (**c**) Boxplot depicting the change in alternative allele frequency of the miRNA SNPs with the top most contribution to linear clustering (N = 27) when comparing goats with high and low genomic ROH content.

To determine which SNPs were contributing the most to the observed differential PCA clustering between “*high*” and “*low*” ROH groups, we performed a variance maximization analysis computing weighted scores over the first two principal component eigenvectors. This allowed us to estimate the importance of each miRNA SNP defining the linear clustering patterns observed after PCA. Among the top most contributing variant sites to variance maximization (N = 27, selected applying a two-component Gaussian mixture approach; see Methods), 21 (77.77%) were located within the pri-miRNA ± 30 bp flanking regions, but just one of them (NW_017189951.1:78275 downstream chi-miR-92b pre-miRNA sequence) was predicted to disrupt a CNNC processing motif (Additional file 12: **Table S12**). Moreover, five SNPs (18.52%) were located in either the stem or the loop of pre-miRNA loci, and one of them was predicted to introduce a novel UGU motif within the loop of chi-miR-100. Interestingly, one miRNA SNP within the seed of chi-mir-425-3p (22:51044384, rs666718867) was also classified among the top most contributing ones to the PCA clustering (Additional file 12: **Table S12**).

We then investigated which goats harbored these miRNA SNPs contributing the most to PCA differential clustering between goats with high and low ROH content. To compare breeds with different sample sizes, we first grouped them according to their geographic origin (see Methods). We then computed weighted standardized allele frequencies according to the number of goats included in each group. Overall, miRNA SNPs were more prevalent in the goat groups with high genomic ROH content (i.e. they had increased alternative allele frequencies), and each geographic group seemed to harbor region-specific polymorphisms (Additional file 13: **Fig. S1**). This might be explained by the overall low alternative allele frequency detected for miRNA SNPs (Additional file 4: **Table S4**, and **Fig. 2a**), meaning that most SNPs are only segregating in few closely related individuals (in terms of geographic or breed origin, see Additional file 14: **Table S13**).

Weighted allele frequencies showed that domestic goats belonging to North Europe (Landrace and Old Irish), Malagasy (Diana, Menabe and Sofia) and Pakistani (Bari, Kachan, Kamori and Pateri) geographic groups harbored a higher number of segregating miRNA SNPs with elevated alternative alle frequencies compared to other goat groups with high or low genomic ROH content (Additional file 14: **Table S13**, and Additional file 13: **Fig. S1**). This pattern was more evident when we just considered the top most contributing polymorphic sites to PCA clustering (N = 27, **Fig. 4b**). Among these, it is worth mentioning 22:51044384 (rs666718867) located within chi-miR-425-3p seed region (alternative allele frequency = 0.426%, see Additional files 4 and 6: **Tables S4**, and **S6**), and segregating in two Kamori goats in a heterozygous state. North European and Pakistani goats from the high ROH group showed a larger pool of segregating miRNA polymorphisms among the top most contributing ones to PCA clustering, but for different sets of SNPs (**Fig. 4b**, Additional file 15: **Table S14**), which is in agreement with the overall regional and breed-specific pattern observed before (Additional file 13: **Fig. S1**). As seen for the 126 miRNA SNPs segregating in goats with high or low ROH contents, 22 out of the 27 top most contributing miRNA SNPs to PCA clustering (81.48%) showed increased alternative allele frequencies in goats from breeds with high ROH content (**Fig. 4c**, Additional file 12: **Table S12**), and this was deemed as a statistically significant enrichment of alternative allele frequencies showing a positive change between goat breeds with high genomic ROHs compared to those with low ROHs (*P*-value = 0.0272). Among the five miRNA SNPs that showed the opposite effect, i.e. increased alternative allele frequency in goats from breeds with low ROHs (Additional file 12: **Table S12**), four showed overall allele frequencies above 10% and were located within the pri-miRNA flanks (± 30 bp of the pre-miRNA hairpin) without altering any processing motifs. The remaining one was located within the apical loop of chi-mir-154a (Additional file 12: **Table S12**).

## Discussion

### Purifying selection constrains variation in functionally relevant miRNA regions

The analysis of segregation patterns of miRNA SNPs across domestic goats worldwide revealed the prevalence of low to extremely low alternative allele frequencies (**Fig. 2a**), with most miRNA loci harboring just one and up to three variants within their sequence. Interestingly, the distribution of such polymorphisms was strongly shaped by functional constraints. In this way, the seed region showed a significantly lower SNP density compared to other miRNA regions or to flanking nucleotides upstream and downstream miRNA loci (**Fig. 2b**). This agrees with the patterns of miRNA variation reported in porcine breeds [29], as well as with previous reports describing an uneven distribution of miRNA variant sites with extremely reduced frequencies in the human genome [30,31], possibly linked to population-specific disease risk [32]. These findings evidence that certain miRNA regions (such as the seed) are subjected to strong purifying selection removing mutations that might alter miRNA binding affinity, as well as miRNA stability or biogenesis [1]. In line with this, miRNAs are among the most evolutionarily conserved loci across vertebrates [33,34].

Within the mature miRNA sequences, miRNA SNPs outside the seed were also found in the anchor (1^st^ 5’ nt) and supplementary pairing (13^th^ to 18^th^ nt) regions. The anchor region is known to play an active role in stabilizing the miRNA loading to the AGO MID domain, which is an essential component to form the active miRISC complex directing miRNA binding activity towards the 3’-UTR of target mRNAs [1,35]. The supplementary pairing region of miRNAs, in contrast, improves the specificity of miRNA binding by compensating seed mismatches typical of metazoan miRNA-mRNA interactions [36], albeit at reduced levels [37]. Other additional regions in the mature miRNA sequence, such as 9^th^ to 12^th^, and 19^th^ to 21^st^-23^rd^ nucleotides, contained more SNPs comparatively (Additional file 4: **Table S4**). This is expected since these regions do not play active roles in neither determining the binding properties of miRNAs nor in the structural configuration of the functional miRISC complex composed by the mature miRNA transcript, the AGO protein, and other effector subunits [1,30].

Beyond the mature miRNA, several SNPs were found in the pre-miRNA stem and the apical loop, some of which may disrupt or generate UGU motifs (Additional file 4: **Table S4**). The UGU motif is a well-known sequence feature sometimes found in the beginning of the pre-miRNA apical loop. This motif helps the DGCR8 dimer jointly with Drosha protein to properly determine the cleavage of the pri-miRNA precursor to generate pre-miRNA hairpins during miRNA maturation [38,39]. Therefore, polymorphisms contributing to alter UGU motifs might modify the efficacy of Drosha processing of pri-miRNA transcripts. Polymorphisms within the apical loop, not necessarily affecting processing motifs, might also have structural consequences on the stability of the pre-miRNA hairpin, thus modifying the accessibility of microprocessor complex substrates [40]. Variant sites within pre-miRNA stems could also alter their hairpin structure, contributing to differences in mature miRNA processing, thus leading to changes in the amount of miRNA molecules able to form active miRISC complexes binding and degrading target mRNA transcripts [41,42].

Moreover, among the set of 134 miRNA SNPs identified within the 30 bp upstream and downstream the pre-miRNA boundaries (Additional file 4: **Table S4**), 18 of them (13.43%) were predicted to create or disrupt processing motifs determining the efficiency of pre-miRNA slicing for miRNA maturation. These motifs (CNNC, UG and GHG), as described for the UGU motif in the pre-miRNA apical loop, are key for a proper recognition of Drosha-DGCR8 protein complex [43,44]. Therefore, any polymorphism altering their sequences could have functional consequences modifying the cleavage of miRNA hairpins during their processing to mature miRNAs, and ultimately impacting their abundance as active post-transcriptional regulators.

### Worldwide domestic goat distribution reflects miRNA polymorphism segregation patterns

By analyzing the reduced set of miRNA SNPs found in our work (N = 206), we were able to detect a clear clustering associated with continental origin (**Fig. 3a**). This agrees well with population structure described in previous studies [14,16,45], and with miRNA-centered analyses using domestic pig breeds [29]. Among African goats, one clearly divergent cluster was identified (**Fig. 3b**), corresponding to Malagasy goats (Additional file 8: **Table S8**). The genetic isolation of Malagasy goats from other breeds in the African continent has been reported before [11,12,45], with a strong founder effect followed by prolonged isolation with no gene flow from other African breeds proposed as the cause for their low genetic diversity [11]. Similarly, two clearly differentiated clusters were found when analyzing the miRNA variability of European domestic goats with Norwegian and Landrace breeds, showing divergent patterns of miRNA polymorphism segregation compared to the remaining European breeds (**Fig. 2c**). The observed divergence of some Norwegian goats might be explained by a founder effect and their continued geographical isolation from other goats with European ancestry, as they were sampled from a population located in Tanzania that was initially established with a few dozen purebred Norwegian Landrace goats in the 80s [46] (Additional file 2: **Table S2**). For Landrace goats, our results also agree with a prolonged genetic isolation described by Cardoso et al. for Finish Landrace goats [11], as well as with recent high levels of inbreeding, founder effect and genetic isolation of Dutch Landrace goats [11,47].

Most of the miRNA SNPs showing relatively elevated alternative allele frequencies (>5%) were found as shared among all African, European and Asian goat populations, while SNPs with extremely reduced frequencies were confined to specific goat breeds or happened in as few as one single individual. This might be a reflection of miRNA SNPs with elevated frequencies being under no selective pressure or even under positive selection [4], and that might have already been segregating in wild or early-domesticated goat populations prior to their worldwide expansion and diversification. In contrast, most of the extremely rare miRNA SNPs found in regions under strict selective pressures might have appeared recently as a product of ineffective purifying selection in local goat populations with either high inbreeding, low population census and/or subjected to genetic drift and founder effects due to prolonged geographical isolation, although additional mechanisms might be at play.

### miRNA polymorphisms mapping to functional regions are more abundant in highly homozygous goat breeds

PCA clustering based on miRNA variation patterns of goat breeds with high and low ROHs revealed three distinct groups with low genetic diversity. Among breeds of European ancestry, Landrace and Old Irish goats clustered apart from other breeds with lower homozygosity (**Fig. 4a**). Contrary to Landrace goats, which increased their census during the second half of the 20^th^ century [47], the Old Irish breed currently has an extremely small population size, with few remaining feral specimens subjected to indiscriminate culling that has led to their near extinction [48]. Their reduced census and overall genetic isolation might have favored the segregation of mutations mapping to functional miRNA regions that might be deleterious. For instance, a SNP located in the seed of chi-miR-1197-3p (21:65190342) was detected in two out of the four Old Irish goats analyzed here, albeit only in a heterozygous state (Additional files 6 and 15: **Tables S6**, and **S14**). Changes in the expression of chi-miR-1197-3p have been linked to whitening of renal brown adipose tissue after birth in goats [49]. Similarly, Malagasy goats, all identified as harboring a high genomic ROH content [12], clustered apart from the other African breeds based on miRNA variation, a pattern that has also been observed at the genome-wide level [45]. An additional third cluster was identified in Asian goats from Pakistan belonging to Pateri, Bari, Kamori, and Kachan breeds, which all show moderate to high levels of inbreeding and low genetic diversity [12,50]. Interestingly, another miRNA SNP detected in the seed region (rs666718867, located in the seed of chi-miR-425-3p) was found segregating in four goats in a heterozygous state, with two of these goats belonging to the Kamori breed, while the other two had unknown breed identification but were sampled in Iran (Additional file 6: **Table S6**). Dysregulation of miR-425-3p expression has been linked to alterations in preadipocyte differentiation, lipolysis and browning of white adipocytes [51].

Despite the putative regulatory and functional impact of SNPs located in the seed of mature miRNAs, their biological relevance is strongly dependent on the developmental context, tissue specificity and metabolic state in which the expression of their hosting miRNAs is elicited. Therefore, any potential regulatory changes shaped by the presence of miRNA SNPs should be taken with caution and put in relation to the expression dynamics of the miRNAs that harbor them. Upon analysis of the miRNA SNPs that contributed the most to the observed divergent clustering between goat breeds with high and low genomic ROH content, the majority of them showed increased alternative allele frequencies in the “high” group, with North European (Landrace and Old Irish breeds), Pakistani, and Malagasy goats showing population-specific segregating miRNA SNPs. Landrace and Malagasy goats have probably been subjected to prolonged geographic isolation and show signs of having undergone bottleneck events [11]. Besides, Pakistani goats— at least those belonging to Bari, Kamori, Kachan, and Pateri breeds—probably reproduce a similar pattern of continued reproductive isolation leading to inbreeding [50,52], as well as specific genomic adaptations to local climates [53], while Old Irish feral goats have an extremely reduced census [48]. This might explain that, while all of them showed distinctive segregating miRNA SNPs, only Northern European and Pakistani goats harbored SNPs mapping to regions with critical functions. Among the few ones that had higher alternative allele frequencies within goats in the “low” group (Additional file 12: **Table S12**), most of them were found in more than 10% of the analyzed goats, possibly indicating they have been segregating for longer under positive selection or with no strong selective pressure.

It is important to note, however, that an elevated genomic ROH content can reflect both recent and ancient inbreeding. In this way, ancient sustained genetic isolation and low effective population sizes lead to increased ROH content genome-wide found in mostly short to middle-sized stretches, while recent population bottlenecks and founder effects leading to higher inbreeding coefficients will produce longer ROHs across the genome (>8–16 Mb) [54,55]. Relatively recent domestication events and breed formation lead to bottlenecks reducing the effective population size, and favoring artificial selection, genetic drift and genetic hitchhiking, i.e. changes in the allele frequency of polymorphic sites segregating together with other variants under positive selection. Moreover, recent bottlenecked populations tend to have a higher proportion of deleterious mutations compared to populations that have been kept at small sizes for longer [17]. All these convergent and at times contradicting genetic changes often result in larger genetic load and a reduced efficacy of purifying selection to remove potentially harmful mutations [17]. In our study, we used average genome-wide ROH content estimates in domestic goat breeds as reported by Bertolini et al. [12], therefore our findings should be put in the context of overall increased homozygosity genome-wide, with no distinction between ancient and modern genetic changes. Nevertheless, the presence of miRNA SNPs potentially disrupting miRNA expression and binding affinities might be a consequence of impaired purifying selection occurring in certain domestic goat breeds with low census and high inbreeding.

## Conclusions

We conclude that goat miRNA genes display very low levels of variation due to the action of purifying selection, particularly in functionally constrained regions such as the seed and the mature miRNA sequence. We have also shown that highly inbred goat breeds with low population size and a high genomic content of runs of homozygosity harbor miRNA SNPs with potentially deleterious effects. Such findings might be explained by the accumulation of deleterious mutations due a reduced efficacy of purifying selection in removing harmful mutations in populations that have undergone continued geographic isolation and relatively recent bottlenecks leading to high inbreeding levels.

## Supplementary Information

**Additional file 1: Table S1.** Annotation of the pre-miRNA (N = 262) and mature miRNA (N = 427) genes in the domestic goat genome assembly (ARS1).

**Additional file 2: Table S2.** Metadata of the 770 domestic goats (*Capra hircus*) from the VarGoats project used in this study.

**Additional file 3: Table S3.** Summary of sampling size, ROH estimates and group classification used for comparing goat breeds with High or Low runs of homozygosity (ROHs) according to Bertolini et al. (2018).

**Additional file 4: Table S4.** Polymorphisms segregating within and around miRNA loci (30 nucleotides upstream and downstream the pre-miRNA boundaries) in domestic goats (N = 770).

**Additional file 5: Table S5.** miRNA polymorphisms showing shared and private segregation in African, Asian, and European domestic goat populations.

**Additional file 6: Table S6.** miRNA polymorphisms located in the seed region and the domestic goat samples where they were found segregating.

**Additional file 7: Table S7.** Principal Component Analysis of the 770 domestic goats included in this study considering segregating miRNA polymorphisms (N = 206).

**Additional file 8: Table S8.** Principal Component Analysis of the African, European and Asian domestic goats included in this study and considering segregating miRNA polymorphisms within each corresponding population.

**Additional file 9: Table S9.** Genotypes of miRNA polymorphisms (N = 206) from all domestic goats used in this study (N = 770).

**Additional file 10: Table S10.** Genotypes of miRNA polymorphisms (N = 126) from domestic goats with High or Low runs of homozygosity (ROHs) according to Bertolini et al. (2018).

**Additional file 11: Table S11.** Principal Component Analysis of domestic goats from breeds with High or Low runs of homozygosity (ROHs) according to Bertolini et al. (2018).

**Additional file 12: Table S12.** Variance maximization weighted scores of miRNA polymorphisms segregating in domestic goat breeds with High or Low runs of homozygosity (ROHs). The top most contributing polymorphic sites are highlighted in grey.

**Additional file 13: Figure S1.** Heatmap representing standardized weighted alternative allele frequencies (*wFrq*) of miRNA polymorphisms segregating in goats with high or low genomic ROH content. Polymorphic sites with standardized wFrq values (in red) are predominantly segregating in the corresponding goat breed.

**Additional file 14: Table S13.** Weighted standardized alternative allele frequencies (Z-scored *wFrq*) of miRNA polymorphisms (N = 126) segregating in domestic goats with High or Low genomic ROHs grouped geographically.

**Additional file 15: Table S14.** Genotypes of the top most contributing miRNA polymorphisms (N = 27) by variance maximization from domestic goats belonging to breeds with High or Low genomic ROHs (N = 151). 0/0: Homozygous genotype for reference allele; 0/1: Heterozygous genotype; 1/1: Homozygous genotype for the alternative allele;./.: Missing data.

## Acknowledgements & Funding

We are grateful to France Génomique “Call for high impact projects” (ANR-10-INBS-09-08). APIS-GENE funded some WGS sequences through ACTIVEGOAT & CAPRISNP projects. We thank the Animal Genetics Division of the French National Institute for Agriculture, Food and Environment (INRAE-GA) for funding VarGoats2 grant, which allowed DNA extraction and genotyping of 384 animals and CRB-Anim, Grant Agreement ANR-11-INBS-0003, (https://crb-anim.fr/) for funding French local breeds sampling. Whole-genome sequencing libraries for the African goats were prepared and sequenced by Edinburgh Genomics and funded via Biotechnology and Biological Sciences Research Council research grant (BBS/OS/GC/000012F) awarded to The Roslin Institute. USDA-ARS with funding from USAID Part of this research was supported by grant PID2022-136834OB-I00 funded by both MCIIN/AEI/10.13039/501100011033 and by “ERDF/EU”. This research was also supported by the SGR Program (2021 SGR 01176) funded by the Direcció General de Recerca (DGR) del Departament de Recerca i Universitats (REU) and by grants SEV-2015-0533 and CEX2019-000902-S funded by MICIU/AEI/10.13039/501100011033, and by the CERCA Programme/Generalitat de Catalunya. Maria Luigi-Sierra was the recipient of a predoctoral fellowship BES-2017-079709 funded by MICIU/AEI/10.13039/501100011033 and by “ESF Investing in your future”. Emilio Mármol-Sánchez also acknowledges financial support from the Villum Fonden (Villum Experiment project no. 57875).

## Authors’ contributions

MA and EM-S conceived the study; the VarGoats consortium provided animal samples and genotype data; the VarGoats consortium curated WGS-data; EM-S performed additional data curation; EM-S analyzed the data. MGLS contributed to data analysis; MA supervised the work; EM-S and MA wrote the original draft. All authors contributed to editing and finalizing the manuscript. All authors read and approved the final manuscript.

## Declarations

### Ethics approval and consent to participate

Blood collection and ear-tag sampling were carried out in accordance with the national regulations from the countries where such samples were collected. In the case of samples sequenced at Génoscope (Evry, France), DNA was imported into France either with authorization 31 555 50, delivered on May 24, 2016, for European countries or covered by an import permit from the DDPP (Direction Départementale de la Protection des Populations) for non-European countries. The only exception was the African animals for which sequencing was performed at Edinburgh Genomicsand the DNA for these libraries was imported into Scotland by permission of the Scottish Government Animal Health and Welfare Division and the UK under generic license IMP-GEN-2008-03.

### Consent for publication

Not applicable

## Data access & availability

The VarGoats dataset is publicly available in the European Nucleotide Archive (ENA) as project number PRJEB37507, which includes fastq and sample description data for 266 animals under accession PRJEB31857, 337 animals under accession PRJEB37122, 29 animals under accession PRJEB37276 and 20 animals under PRJEB37208. A total of 290 additional sequences were retrieved from public databases and 217 from NextGen Consortium projects (PRJEB3134, PRJEB3135, PRJEB4371, PRJEB5166, PRJEB3136 and PRJEB5900 studies). Individual accession numbers are listed in Additional file 2: **Table S2**.

Use of these data is regulated by a data sharing agreement which is available here: http://www.goatgenome.org/vargoats_agreement.html. This agreement states that it is mandatory to contact the VarGoats steering committee to discuss the utilization and inclusion of data generated by the VarGoats Consortium in any present or future publication. No publications can be generated from the VarGoats dataset until the main papers derived from this project are published in scientific journals. Scientists who have signed the agreement have access to 30 VCF files split by chromosome for SNPs and an additional VCF for InDels. They can also access the VCF used for the truth sets and the VQSR VCF file for SNPs. The variant data for this study have been deposited in the European Variation Archive (EVA) at EMBL-EBI under accession number PRJEB90141. Analyses: ERZ26880186. (https://www.ebi.ac.uk/eva/?eva-study=PRJEB90141).

---

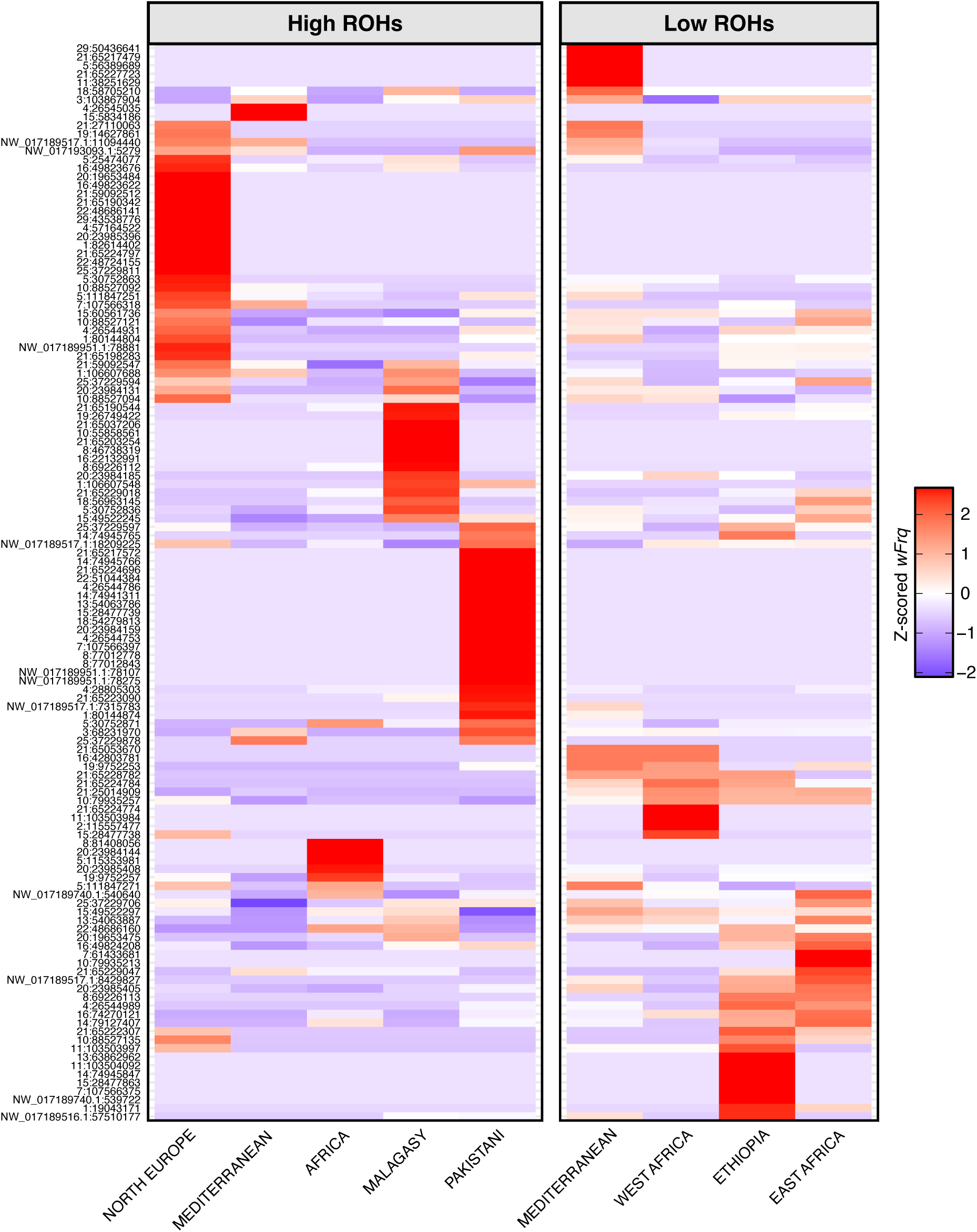

